# Predicting the Reprogrammability of Human Cells Based on Transcriptome Data and SGD Classifier with Elastic-Net Regularization

**DOI:** 10.1101/2022.07.18.500480

**Authors:** Gorkem Saygili, Mine Turktas, Cansu Gurcan, Lucia Gemma Delogu, Acelya Yilmazer Aktuna

## Abstract

Cell reprogramming has shown considerable importance in recent years; however, the programmability of cells and efficiency of reprogramming varies across different cell types. Considering several weeks of cell programming process and costly programming agents used through the process, every failure in reprogramming comes with a significant burden. Better planning for reprogramming experiments could be possible if there is a way of predicting the outcome of reprogramming before the experiments using transcriptome data. In this study, we have accessed the transcriptome data of successful or unsuccessful programming studies published in literature and constructed a Stochastic Gradient Descent (SGD) classifier with Elastic-Net regularization for predicting whether the cell lines are reprogrammable. We tested our classifier using 10-fold cross validation over cell lines and on each cell separately. Our results showed that it is possible to predict the outcome of cell reprogramming with accuracies up to 98% and Area Under the Curve (AUC) scores up to 0.98%. Considering the success of our experimental outcomes we conclude that an outcome of a cell reprogramming experiment can be predicted with high accuracy using machine learning on transcriptome data.

## Introduction

There is an increasing trend for using artificial intelligence (AI) in healthcare and biomedical sciences. As machine learning (ML) algorithms and computing infrastructure develop, these technologies become part of diagnosis and treatment of various human pathologies. Considering the importance of applying personalized medicine and the growing trend on the application of ML techniques in medicine field, numerous studies made use of these technologies to achieve cell reprogramming, stem-cellbased therapies and cancer cell prediction^1–5^. There are also studies in literature aimed for precision oncology: cancer sub-typing based on various different genomic, molecular and pathological parameters^6–9^.

It is very well-known that cells can be reprogrammed back to a pluripotent stage by overexpression of reprogramming factors. Cell reprogramming through forced expression of reprogramming factors (including Oct3/4, Klf4, Sox2, c-Myc) has been reported in several studies in literature^10–12^. Different cell types have been successfully reprogrammed until now, including fibroblasts, hematopoietic stem cells, hepatocytes and even various cancer cells. For example, several groups have tried to reprogram cancer cells via different protocols including the use of embryonic microenvironment, nuclear transfer or transfection with the reprogramming factors resulting in altered tumorigenicity^13–18^. Even though cell reprogramming has shown considerable importance in recent years, the programmability of cells and outcome of reprogramming varies across different cell types. Furthermore, cell reprogramming requires several weeks of tedious effort and considerable cost regarding to utilized reprogramming agents and materials. Therefore, it is clear that there is a need for predicting the outcome of reprogramming in this field to help scientists to better plan for the reprogramming experiments. ML algorithms have been used in single-cell RNA sequence analysis specifically for cell identification and clustering^19–23^. Considering the vast variety of different ML algorithms, it is important to choose the best performing algorithm on the applied domain. Abdelaal et al.^22^ compared the performances of 22 classification methods including the most popular ones with Random Forest (RF), Support Vector Machines (SVM), and Neural Networks on 27 publicly available single cell RNA sequencing datasets and revealed that linear SVM had the overall best overall performance. In a different study, Michielsen et al.^23^ compared linear SVM and one-class SVM and found that linear SVM outperformed one-class SVM on cell classification using RNA-seq data. The datasets that were used for both studies contain cells up to hundreds of thousands. Having a large number of samples is crucial since, depending on the their complexity, ML algorithms may require considerable amount of samples for training. Large amount of data are at the disposal of the researchers to train their algorithm to do cell classification however such data is not available for cell reprogramming in particular for the failure of the programming scenarios. When the data is limited, designing an accurate ML algorithm becomes a challenging task because of the well-known overfitting problem^24^. To cope with it, regularization parameters such as L1 and L2 regularization have been used frequently. Elastic-Net is a combination of L1 and L2 regularizations to achive the best possible performance against overfitting. It has been preferred recently for predicting immune cell tags because of its favorable performance on high dimensional data^25^.

There are two main contributions of this study:

- For the first time, we show that programming outcome of a cell can be predicted before the programming process using ML and transcriptome data.
- Elastic-Net regularization with an SGD classifier provides better performance with limited number of samples than the overall best performer of previous studies, linear SVM ^22,23^.

Our second contribution is particularly important when there is limited number of cells such as rare cancer cell populations and the dimensionality of the data is large which is in general true for transcriptome data. Because in such instances, using deep neural networks^26–28^ could not be an option because of their large sample size demand for accurate training.

## Results

### Performance comparison of 10 Machine Learning Algorithms Over the Replicates Indicates Predicting the Outcome of Reprogramming is Achievable with High Accuracy

In order to achieve a higher sample number to establish ML-based algorithms, we accessed transcriptome data of various reprogramming studies from the literature that resulted in either success or failure. Transcriptome reads that have been downloaded included that of melanoma cells from Castro-Perez et al.^29^, human CD34+ cells from Friedli et al.^30^, human fibroblast cells from different studies^31–36^, human acute myeloid leukemia cells from Chao et al.^37^, human thyroid-carcinoma cell-lines from Kong et al.^38^. Out of these 18 cell lines (consisting of 38 replicate (cells)), 12 cell lines (26 replicates) have been reported to be successfully reprogrammed whereas 6 (12 replicates) of them failed to generate iPSC colonies^29–36^. On this data we have attempted to establish ML-based algorithm to predict the programmability of cells based on their transcriptome data of prior to reprogramming.

We predicted whether each a replicate is reprogrammable or not with leave-one-out cross validation (LOOCV) experiment. Through the experiment, a replicate was left out and the remaining replicates were used as the training set. Since our dataset was reasonably small, we aimed to have the maximum number of training samples with this approach. We repeat LOOCV experiment 10 times to minimize the stochastic results as much as possible. The hyperparameter optimization step was done using 3-fold CV on the training set and the prediction was made for the replicate in the test set with the best-performing hyperparameters on the training set. The Receiver Operating Characteristic (ROC) and Precision-Recall curves are presented in Figures 1a and 1b, respectively. From both results, Linear Support Vector Machine (SVM), Generalized Linear Model (GLM), Random Forest (RF), Naive Bayes (NB), Stochastic Gradient Descent with Elastic-Net regularization (SGD), and XGBoost provided top performances with over 0.9 AUC, accuracy, and AP scores. In contrast to these top performers, Two classes of reprogrammable and non-reprogrammable cells were again separable with reasonably high accuracy which can be perceived from the density plots and the ROC curve. The AUC score of 1 indicates almost perfect classification performance. To have a detailed performance comparison, Table 1 presents the accuracy, sensitivity, specificity, precision, F1 score and Matthew’s Correlation Coefficient (MCC) scores of each classifier for LOOCV experiment. In particular, MCC scores are more reliable compared to other measures, thanks to its balanced use of all confusion matrix components^39^. Based on MCC score, SGD with Elastic-Net regularization (SGD_Elastic), NB, GLM, and SVM performed on par and outperformed the other classifiers. All of these four classifiers also achieved high accuracy, sensitivity. specificity, precision and F1-score close, if not equal to 100%. The overall results indicate that the reprogramming outcome of a replicate can be predicted using transcriptome data and ML.

**Table 1.** Accuracy, sensitivity, specificity, precision, F1-score and Matthew’s correlation coefficient (MCC) scores of 9 classifiers from LOOCV experiments. The top performance for each score is identified as bold.

| Classifier | Accuracy | Sensitivity | Specificity | Precision | F1 Score | MCC |
| --- | --- | --- | --- | --- | --- | --- |
| Adaboost | 0.85 | 0.87 | 0.80 | 0.90 | 0.89 | 0.66 |
| KNN | 0.87 | 0.85 | 0.92 | 0.96 | 0.90 | 0.73 |
| LDA | 0.55 | 0.50 | 0.67 | 0.76 | 0.60 | 0.16 |
| SVM | 0.97 | **1.00** | 0.92 | 0.96 | **0.98** | 0.94 |
| GLM | 0.97 | 0.96 | **1.00** | **1.00** | **0.98** | 0.94 |
| NB | 0.97 | **1.00** | 0.92 | 0.96 | **0.98** | 0.94 |
| XGBoost | 0.92 | 0.92 | 0.92 | 0.96 | 0.94 | 0.82 |
| RF | 0.95 | 0.92 | **1.00** | **1.00** | 0.96 | 0.89 |
| SGD_Elastic | **0.98** | 0.98 | 0.98 | 0.99 | **0.98** | **0.95** |

**Figure 1.**
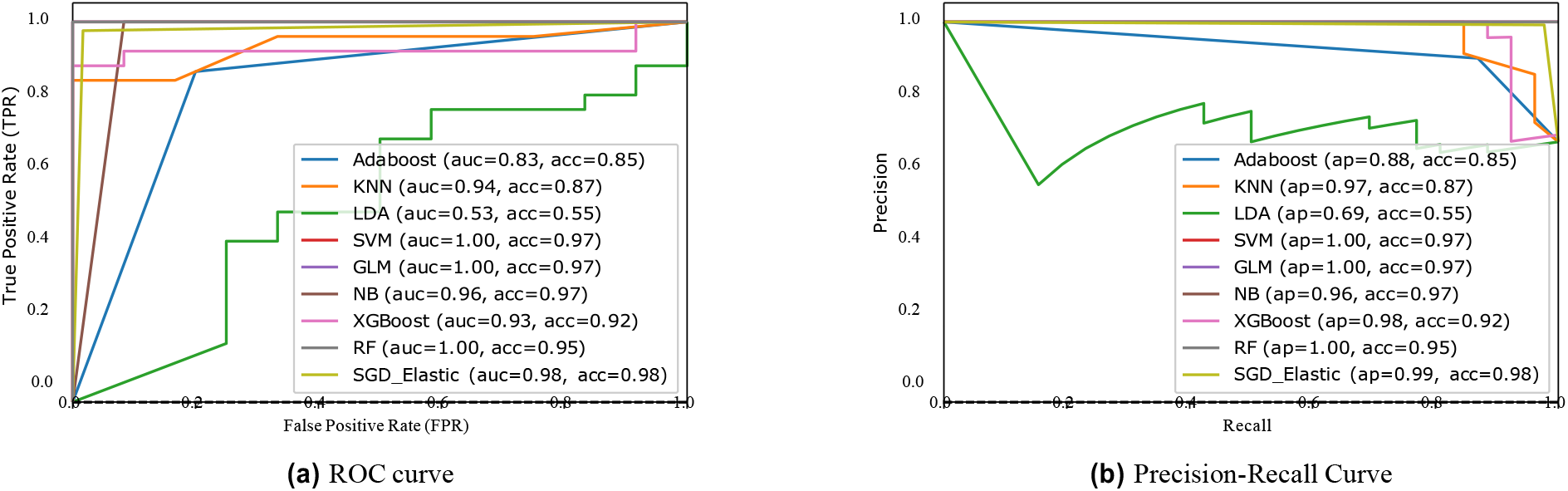
(a) ROC curve and (b) Precision-Recall curves of utilized classifiers from LOOCV experiments. Area Under ROC Curve (AUC), accuracy, and Average Precision score (AP) scores are provided for each classifier.

#### Performance comparison of 10 Machine Learning Algorithms Over the Celllines Predicting the Outcome of Reprogramming is Achievable with High Accuracy

Since ML algorithms may exploit transcriptional similarities between the replicates, we conducted another experiment, so called cell line experiment, in which we separated our test set to contain all the replicates from a cell line and hence there is no replicate from that cell line in the training set. Since there is no replicates of the same cell line in both training set and the test set, the ML algorithms can only exploit transcriptional differences between successfully programmed cells and the unsuccessful ones. We repeated cell line experiment 10 times to minimize the stochastic results as much as possible. Same as the LOOCV experiment, the hyperparameter optimization step was done using 3-fold CV on the training set. The prediction was made all replicates in a cell line in the test set with the best-performing hyperparameters on the training set. The Receiver Operating Characteristic (ROC) and Precision-Recall curves are presented in Figures 2a and 2b, respectively. Different from the previous results, classifiers generally performed less accurately in this experiment. Two ensemble learners, Adaboost and XGBoost, and LDA performed less accurately compared to other classifiers. Table 2 presents the accuracy, sensitivity, specificity, precision,F1 score and MCC scores of each classifier for the cell line experiment. The decrease in the performance of classifiers is clearly observed from the MCC scores. SGD classifier with Elastic-Net regularization had on par performance with LOOCV experiment however all other classifiers performed considerably worse than SGD. Furthermore, all the remaining scores of SGD is also similar to its LOOCV performance, proving that SGD classifier with Elastic-Net regularization achieved the best overall performance in the prediction of reprogramming outcome.

**Table 2.** Accuracy, sensitivity, specificity, precision, F1-score and Matthew’s correlation coefficient (MCC) scores of 9 classifiers from Cellline experiments. The top performance for each score is identified as bold.

| Classifier | Accuracy | Sensitivity | Specificity | Precision | F1 Score | MCC |
| --- | --- | --- | --- | --- | --- | --- |
| Adaboost | 0.88 | 0.91 | 0.83 | 0.92 | 0.91 | 0.73 |
| KNN | 0.89 | 0.88 | 0.92 | 0.96 | 0.92 | 0.77 |
| LDA | 0.68 | 0.58 | 0.92 | 0.94 | 0.71 | 0.46 |
| SVM | 0.92 | **1.00** | 0.75 | 0.90 | 0.95 | 0.82 |
| GLM | 0.89 | 0.85 | **1.00** | **1.00** | 0.92 | 0.80 |
| NB | 0.92 | 0.92 | 0.92 | 0.96 | 0.94 | 0.82 |
| XGBoost | 0.63 | 0.73 | 0.42 | 0.73 | 0.73 | 0.15 |
| RF | 0.89 | 0.85 | **1.00** | **1.00** | 0.92 | 0.80 |
| SGD_Elastic | **0.97** | 0.98 | 0.96 | 0.98 | **0.98** | **0.93** |

**Figure 2.**
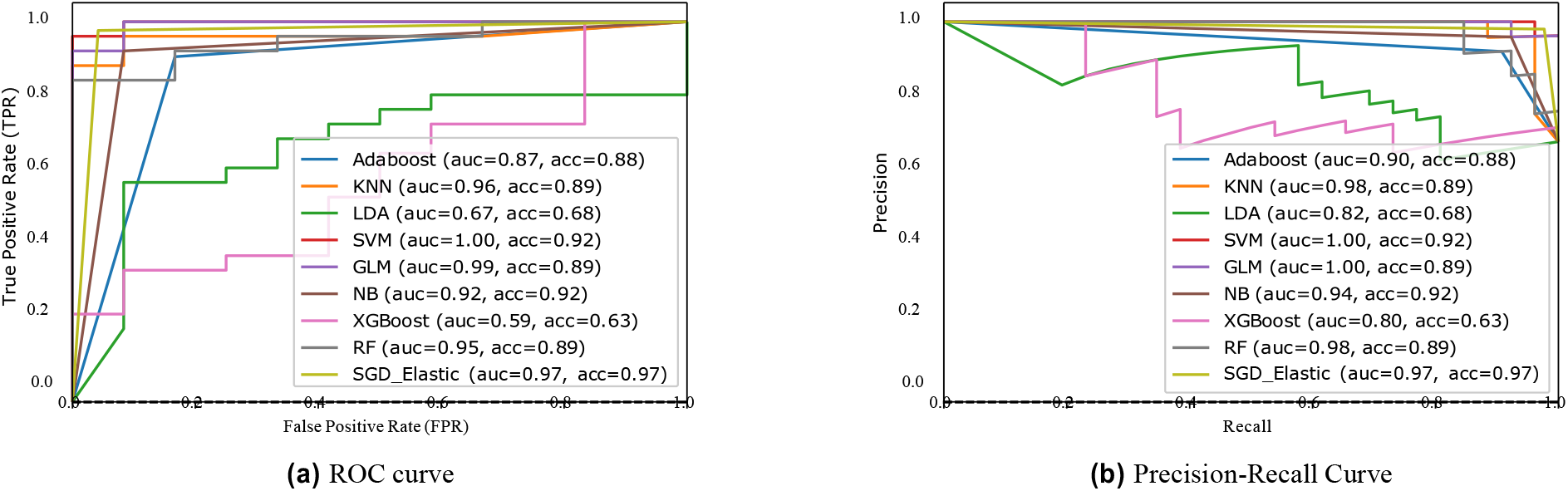
(a) ROC curve and (b) Precision-Recall curves of utilized classifiers on cell-line experiments. AUC, accuracy, and AP scores are provided for each classifier.

#### SGD classifier revealed important genes in the prediction of reprogramming outcome

The weights assigned to features can be used to assess the importance of that feature for the prediction of the target variable. SGD classifier with Elastic-Net regularization provides non-negative feature weights and considerable amount of them can be zero on a high dimensional dataset because of L1 regularization. Hence, SGD classifier intrinsically has feature selection capability and eventually features with larger weights indicate important features for the classification task. Figure 3a and 3b show the five genes that have the highest weights in the classification task. Both figures indicate the same set of genes namely, RN7SK, 7SK, RN7SL2, RN7SL1, and MT-RNR2. The others indicate the sum of the weights of the remaining genes.Since both experiments indicate the same set of genes, we advocate that these genes play a crucial rule for the outcome of reprogramming.

**Figure 3.**
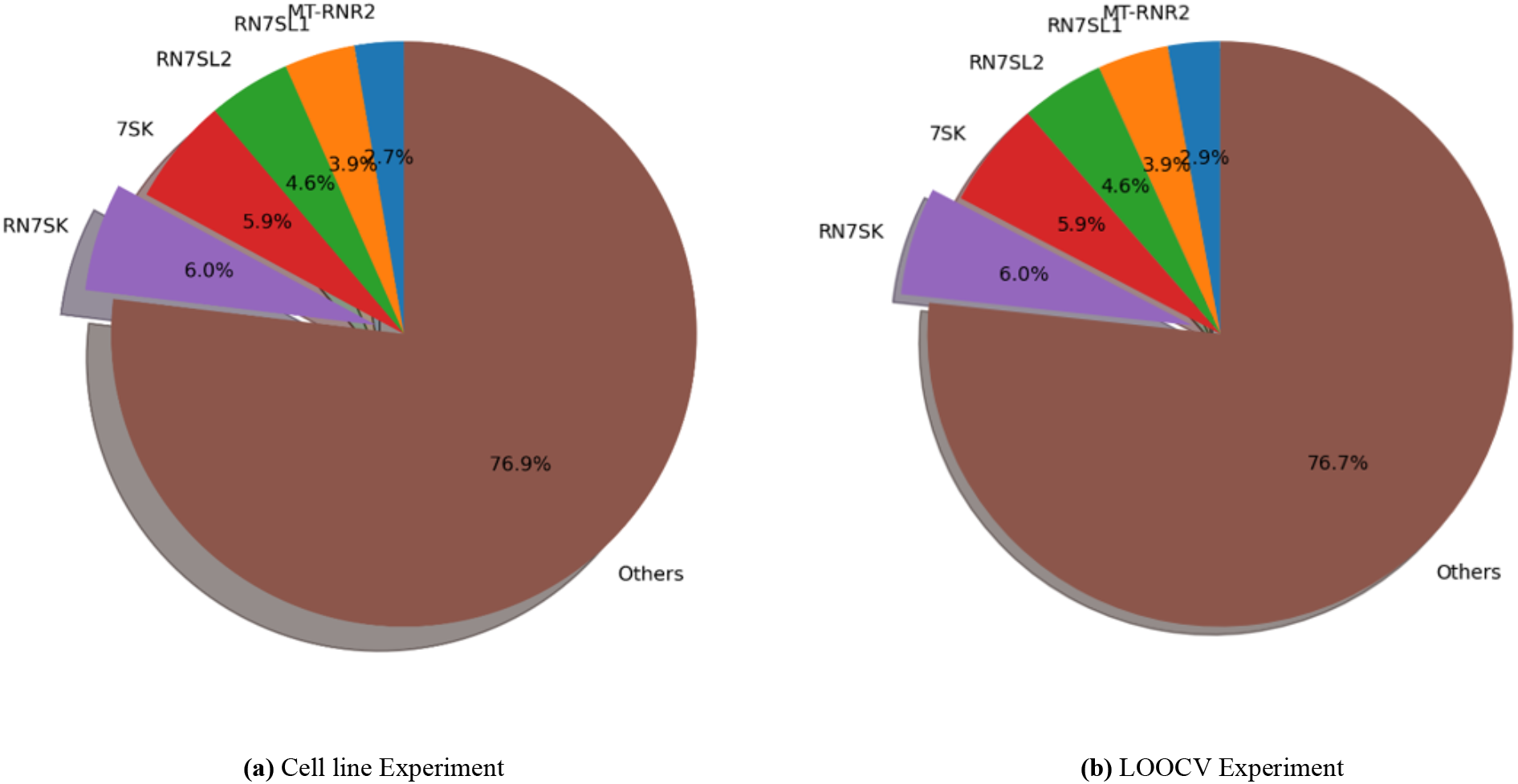
Top five genes that has the highest weights in SGD classifier with ElasticNet regularizationm, indicating their importance on the classification result.

#### Dimensionality reduction methods reveal the distinction between successfully programmed and failed to be reprogrammed replicas

Dimensionality reduction such as Principal Component Analysis (PCA)^40^, t Stochastic Neighbor Embedding (t-SNE)^41^, and Unifold Manifold Approximation and Projection (UMAP)^42^ are commonly used with transcriptome data to visualize the clusters of the samples. Two popular visualization algorithms, t-SNE and UMAP, provide faithful representations of high dimensional data, preserving close neighborhoods in the embeddings. All of the embedding results are presented in Fig. 4 which clearly show that there are two clusters consists of successfully reprogrammed and failed to be reprogrammed replicates. In particular, the two clusters are clearly visible on t-SNE (Fig. 4b) and UMAP (Fig. 4c) embeddings. Two clusters in the lower dimensional space clearly indicates the existing similarities in terms of transcriptome profiles of the in between successfully programmed replicates and in between failed to be programmed replicates.

**Figure 4.**
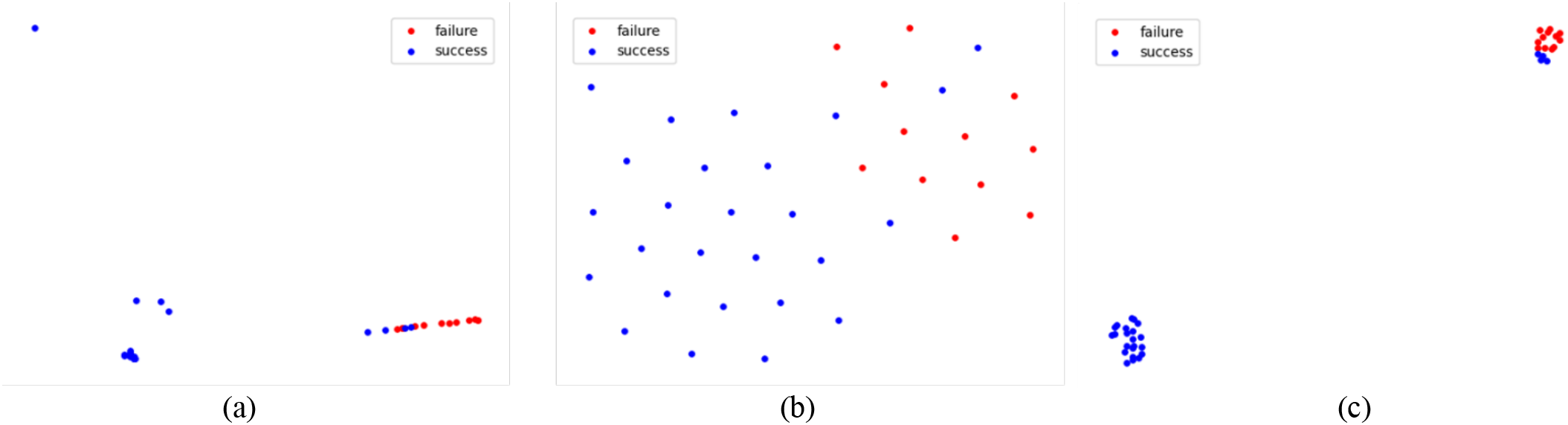
(a) PCA, (b) t-SNE, and (c) UMAP embeddings of the successfully reprogrammed (success) and failed to be reprogrammed (failure) replicates.

## Discussion

Continued research and development in cell reprogramming technology will be a valuable tool for more refined in vitro disease models, identification of disease biomarkers and ultimately for the development of novel treatment protocols for patients in the future^10,14–17^. For example, cancer cell reprogramming studies can provide additional opportunities including: modifying cancer stem cell populations, which would be an important tool to better understand their biological properties in resistant tumors; engineering the resulting reprogrammed cancer cells with altered tumorigenicity in order to develop novel cancer vaccines; using cancer-derived iPSC for pharmacological screenings; or creating a powerful in vitro model to understand the epigenetic and genetic mechanisms taking place during tumor development and progression^10^. Published studies indicate that the starting cell status is highly variable and, in turn, may correspond to partial or complete pluripotent states^14^. Therefore, there is a need for establishing ML-based prediction prior to starting the reprogramming studies. Looking at the recent years, there is a growing trend for using ML-based approaches in biomedical sciences. This study aims to predict the reprogrammability of cells based on a ML-based algorithm which was trained on already published data. It is of crucial importance to validate the predictive performance of these models; therefore, this study makes an important contribution in the field by both establishing a ML-based prediction.

In this study, we aim to predict the reprogrammability of various cell types using 9 different classifiers including the best classifier for single cell classification based on transcriptome data, linear SVM, as mentioned in^22,23^. Both LOOCV and Cell line experiment results showed that accurate predictions up to 97% accuracy are possible using transcriptome data before the reprogramming. Even more importantly, more reliable measures such as F1 score and MCC are also both over 0.93 for both experiments with SGD Classifier with Elastic-Net regularization. These results show that it is possible to predict the outcome of a reprogramming experiment based on transcriptome data and ML. Although we have not tested, we believe that stochastic gradient descent which is the main optimizer of deep neural networks, can be trained and tested faster than SVM.

An important bottleneck of this study is the size of its dataset. Our dataset is limited in terms of number of samples because the transcriptome data with unsuccessful reprogramming outcome is limited. Considering the high dimensionality of transcriptome data and limited number of samples, overfitting problem becomes a crucial concern. The difference between SGD classifier and linear SVM in our case is the Elastic-Net regularization in which Ridge (L2) and Laso (L1) regularizations are combined. In comparison to Linear SVM which incorporates only Ridge regularization, Elastic-Net provides considerable robustness against overfitting and increases the overall performance as depicted in our results. Although plain SGD classifier with Ridge loss is similar to linear SVM, SGD classifier with Elastic-Net regularization clearly outperformed linear SVM. The power of Elastic-Net regularization against overfitting has been also showed in^25^. We believe the addition of Laso regularization which can cause some of the weights to be exact zero in the classifier causes an effect of adaptive feature selection through the training and leads to a considerable increase in performance. This advantage can be exploited in different classification tasks with transcriptome data when the available data is limited in terms of samples.

Since our overall classification performance is high, we explored the genes that played crucial role in the prediction of the reprogramming experiment. Our results show that RN7SK (ENSG00000283293), 7SK (ENSG00000202198), RN7SL2 (ENSG00000274012), RN7SL1 (ENSG00000276168), and MT-RNR2 (ENSG00000210082) were the top five important genes that contributed to the classification. RN7SK has shown to play a role in cellular differentiation^43^ and senescence^44^. Studniarek et al.^45^ have found that 7SK play an important role in proper stress response and cell survival. Perhaps more importantly, Eilebrecht et al.^46^ has shown that 7SK directly affects HMGA1 function which induces the pluripotency gene expression. In their work, Nabet et al.^47^ showed that RN7SL1 activates PRR RIG-I which eventually enhances tumor growth.Following up these studies, our analysis is also suggesting that non-coding RNAs are important regulatory elements in cellular reprogramming and differentiation processes. On the other hand, Palombo et al.^48^ revealed the role of mitochondrial DNA (MtDNA) variants’affect on the phenotype of iPSCs and their healthiness. Therefore, based on our analysis, it would be important to perform variant analysis of key MtDNA genes such as MT-RNR2 prior to start of cellular reprogramming experiments in order to ensure good quality iPSC generation. We believe that analyzing the feature importances of accurate classifiers further could reveal many more essential genes that play a crucial role in explaining various factors that affect the outcome cell reprogramming.

## Methods

### Data retrieval

The transcriptome data of Friedli et al.^30^, Tanaka et al.^31^, Castro-Perez et al.^29^ was used for the analyses and data was retrieved from NCBI Gene Expression Omnibus database. The accession numbers were given in the Supplementary Table 1. Our data consisted of 38 replicates samples from 18 cell lines and 57250 expression levels. Each cell line corresponded to one of our samples whereas each expression level constituted our features. In total, 26 of those cells were reprogrammed successfully (labeled as 1), whereas the remaining 12 were failed to be reprogrammed (labeled as 0).

### RNA-Seq data analysis

The RNA-seq data analysis was performed using the new version of the Tuxedo protocol with HISAT2, StringTie and Ballgow^49^. Homo sapiens GRCh38 reference genome was used for genome indexing. The transcriptome reads were aligned to the indexed genome using Hisat2^50^. Mapped reads of each sample were assembled and the transcript counts in FPKM were obtained using StringTie^49^.

### Application of machine learning algorithms

The majority of 9 classifiers were chosen based on the classifiers that were used in^22^. Multi layer perceptron (MLP) classier has not been used since depending on the number of hidden layers, MLP classifiers need substantial amount of data to be trained successfully. Rather than MLP, different from^22^, we incorporated boosting ensemble learners such as Adaboost and XGBoost. Additionally, we included SGD classifier with Elastic-Net regularization which achieved the highest overall performance.

An important step in machine learning experiments is the hyperparameter optimization step. The hyperparameters differ among different ML algorithms, for example number of estimators and maximum depth are important hyperparameters of RF classifiers whereas the type of the kernel (RBF, linear, etc.), gamma and C values are important hyperparameters of SVM classifiers. To optimize these parameters for our ML algorithm, we applied 3-fold cross validation and grid search on the training set for each of our experiments on training set. Hence, we optimize each ML algorithm on the training set and tested this optimized ML algorithm on the test set to obtain the best possible predictions.

There are several scores to quantify the performance of a classifier. Among those, accuracy is the most common score. Yet, accuracy does not always present the whole performance especially when class labels are imbalanced. Another performance measure is the Receiver Operator Characteristic (ROC) curve and Area Under the Curve (AUC) score measured from the ROC. An AUC score close to 0.5 indicates that the classifier produces random predictions whereas an AUC score close to 1 signifies high performance. In our experiments, we used several classifier performance measures such as F1 Score, AUC, and Matthew’s Correlation Coefficient (MCC) to interpret the performance of our classifier.

We used Python programming language and scikit-learn toolkit (version 0.22) in our experiments on a standard computer with 32 GB of memory.

LaTeX formats citations and references automatically using the bibliography records in your .bib file, which you can edit via the project menu. Use the cite command for an inline citation, e.g.^**?**^.

For data citations of datasets uploaded to e.g. *figshare*, please use the howpublished option in the bib entry to specify the platform and the link, as in the Hao:gidmaps:2014 example in the sample bibliography file.

## Acknowledgements

A.Y. would like to acknowledge the Turkish Academy of Sciences for financial support under the young investigator programme (GEBIP2018).

## Author contributions statement

G.S. conceived the machine learning experiment(s) and analyzed the results, M.T.E. conducted the integration/registration of different datasets, C.G., L.G.D. and A.Y.A find and collected the relevant datasets. All authors reviewed the manuscript.

## Additional information

The authors have no conflicts of interest to declare. All co-authors have seen and agree with the contents of the manuscript and there is no financial interest to report. We certify that the submission is original work and is not under review at any other publication.

